# Reliability and generalizability of neural speech tracking in younger and older adults

**DOI:** 10.1101/2023.07.26.550679

**Authors:** Ryan A. Panela, Francesca Copelli, Björn Herrmann

## Abstract

Neural tracking of continuous, spoken speech is increasingly used to examine how the brain encodes speech and is considered a potential clinical biomarker, for example, for age-related hearing loss. A biomarker must be reliable (intra-class correlation [ICC] >0.7), but the reliability of neural-speech tracking is unclear. In the current study, younger and older adults (different genders) listened to stories in two separate sessions while electroencephalography (EEG) was recorded in order to investigate the reliability and generalizability of neural speech tracking. Neural speech tracking was larger for older compared to younger adults for stories under clear and background noise conditions, consistent with a loss of inhibition in the aged auditory system. For both age groups, reliability for neural speech tracking was lower than the reliability of neural responses to noise bursts (ICC >0.8), which we used as a benchmark for maximum reliability. The reliability of neural speech tracking was moderate (ICC ∼0.5-0.75) but tended to be lower for younger adults when speech was presented in noise. Neural speech tracking also generalized moderately across different stories (ICC ∼0.5-0.6), which appeared greatest for audiobook-like stories spoken by the same person. This indicates that a variety of stories could possibly be used for clinical assessments. Overall, the current data provide results critical for the development of a biomarker of speech processing, but also suggest that further work is needed to increase the reliability of the neural-tracking response to meet clinical standards.

**Significance statement:** Neural speech tracking approaches are increasingly used in research and considered a biomarker for impaired speech processing. A biomarker needs to be reliable, but the reliability of neural speech tracking is unclear. The current study shows in younger and older adults that the neural-tracking response is moderately reliable (ICC ∼0.5-0.75), although more variable in younger adults, and that the tracking response also moderately generalize across different stories (ICC ∼0.5-0.6), especially for audiobook-like stories spoken by the same person. The current data provide results critical for the development of a biomarker of speech processing, but also suggest that further work is needed to increase the reliability of the neural-tracking response to meet clinical standards.

## Introduction

Understanding how speech is processed in the brain is important for clinical applications, such as age-related hearing loss, dementia, and stroke (Schneider et al., 2002; Tyler and Marslen-Wilson, 2008; Olichney et al., 2011). Traditional approaches to understanding speech processing have relied on word or sentence materials presented in random, disconnected order (Friederici et al., 1993; Marinkovic et al., 2003; Herrmann et al., 2011). Such materials lack a topical thread, are not very interesting to the listener, and thus may not capture how speech is processed in real life (Hamilton and Huth, 2020). Over the past decade, experimental and analytic approaches have substantially advanced to capture speech processing for more naturalistic, continuous speech, such as spoken stories (Ding and Simon, 2012; Crosse et al., 2016; Brodbeck and Simon, 2020; Hamilton and Huth, 2020; Herrmann and Johnsrude, 2020).

Possibly the most used approach to investigate the extent to which continuous, spoken speech is encoded in the brain is the temporal response function and related encoding/decoding models that, for example, predict electroencephalography/magnetoencephalography (EEG/MEG) activity from speech features (Lalor and Foxe, 2010; Crosse et al., 2016; Brodbeck and Simon, 2020; Crosse et al., 2021). Quantifying how well the brain tracks a specific speech feature provides an estimate of how well the feature is encoded. Especially the neural tracking of the low-frequency (<10 Hz) acoustic amplitude envelope of speech has been extensively studied, for example, in the context of selective attention (Fiedler et al., 2019; Brodbeck and Simon, 2020; Emily et al., 2022) and speech masked by background sound (Schmitt et al., 2022; Synigal et al., 2023; Yasmin et al., 2023). Investigating the neural tracking of the speech envelope is a valuable approach, because the envelope is important for speech intelligibility (Shannon et al., 1995) and envelope tracking predicts speech intelligibility to some extent (Ding et al., 2014; Vanthornhout et al., 2018; Lesenfants et al., 2019; see discussion in Gillis et al., 2022). Moreover, calculating the envelope is easy and available in various toolboxes (Crosse et al., 2016; Crosse et al., 2021), whereas other recent approaches are more complex (Broderick et al., 2018; Broderick et al., 2021).

Neural speech-tracking approaches are increasingly used to understand clinical phenomena, such as age-related hearing loss (Presacco et al., 2019; Decruy et al., 2020a; Schmitt et al., 2022) and stroke-related or dementia-related aphasias (Dial et al., 2021; Kries et al., 2023). For example, aging and hearing loss are associated with enhanced neural tracking of the speech envelope (Presacco et al., 2016b; Decruy et al., 2020a), which may be the result of a loss of cortical inhibition resulting from periphery deafferentation (Caspary et al., 2008; Herrmann and Butler, 2021a). Given the success of the speech-tracking approach, researchers have suggested that the neural-tracking response could be used as a biomarker for speech-processing pathologies (Gillis et al., 2022; Palana et al., 2022; Schmitt et al., 2022). Yet, for the neural-tracking response to be a useful as a biomarker it must be reliable, but its reliability is currently unclear.

Reliability is investigated by conducting the same test procedures twice (Lockhart, 1998). The intra-class correlation (ICC; Shrout and Fleiss, 1979; McGraw and Wong, 1996; Koo and Li, 2016) – a metric that captures both the degree of correlation and agreement between two measurements – may be used to assess reliability, although some previous EEG/MEG research has used Spearman’s or Pearson’s correlation instead, which only assesses correlation (Tervaniemi et al., 1999; McEvoy et al., 2000; Cabral-Calderin and Henry, 2022). Neural responses to tones, noises, and vowels have moderate to high reliability (Tervaniemi et al., 1999; Legget et al., 2017; Bidelman et al., 2018), whereas little is known about the reliability of the neural tracking of continuous speech. Moreover, in clinical contexts, individuals would ideally listen to unique stories – for example, between appointments – to avoid influences of prior knowledge on the neural response. Hence, it would be beneficial if the neural-tracking response generalized across different stories.

The current study provides an extensive account of the reliability and generalizability of the neural-tracking response across different speakers, stories, and noise conditions in younger and older adults. The data are critical for research aiming to use the neural-tracking response as a biomarker for the assessment of auditory or cognitive impairments.

## Methods and materials

### Participants

Twenty-two younger adults (median: 22 years; range: 19–34 years) and 22 older adults (median: 70.5 years; range: 56-77 years) participated in the current study. Sixteen younger adults identified as female or women, five as male, and one as non-binary. Fourteen older adults identified as female and eight as male. Twelve younger and 20 older adults identified as native English speaker, whereas the other participants were highly proficient English speakers. Participants who indicated having a non-English first language, nevertheless, grew up in English-speaking countries (mostly Canada) and have been speaking English since early childhood (<5 years of age). Participants reported having no neurological disease, except for one older adult who indicated having trigeminal neuralgia (not affecting participation and results). Participants reported having normal hearing abilities. Participants did not wear hearing aids nor were they prescribed one.

Each participant took part in two sessions, separated by at least one week (median: 13 days; range: 7-59 days). Participants gave written informed consent prior to the experiment and were paid $7.50 CAD per half-hour for their participation. The study was conducted in accordance with the Declaration of Helsinki, the Canadian Tri-Council Policy Statement on Ethical Conduct for Research Involving Humans (TCPS2-2014), and was approved by the Research Ethics Board of the Rotman Research Institute at Baycrest.

### Sound environment and stimulus presentation

Data collection was carried out in a sound-attenuating booth. Sounds were presented via Sennheiser (HD 25-SP II) headphones and a RME Fireface 400 external sound card. Stimulation was run using Psychtoolbox in MATLAB (v3.0.14; MathWorks Inc.) on a Lenovo T480 laptop with Microsoft Windows 7. Visual stimulation was projected into the sound booth via screen mirroring. All sounds were presented at about 75 dB SPL.

### Hearing assessment

For each participant, audiometric thresholds were estimated for pure tones at frequencies of 0.25, 0.5, 1, 2, 4, and 8 kHz (Figure 1A). Mean thresholds averaged across 0.25, 0.5, 1, 2, and 4 kHz were higher for older compared to younger adults (t_42_ = 5.804, p = 7.6 · 10^-7^, d = 1.75; Figure 1). Although these thresholds are mostly clinically ‘normal’ for age according to the ISO-7029 standard (https://www.iso.org/standard/42916.html), elevated thresholds are consistent with the presence of mild-to-moderate hearing loss in the current sample of older adults, as would be expected (Moore, 2007; Plack, 2014; Presacco et al., 2016b; Herrmann et al., 2018, 2022).

**Figure 1:**
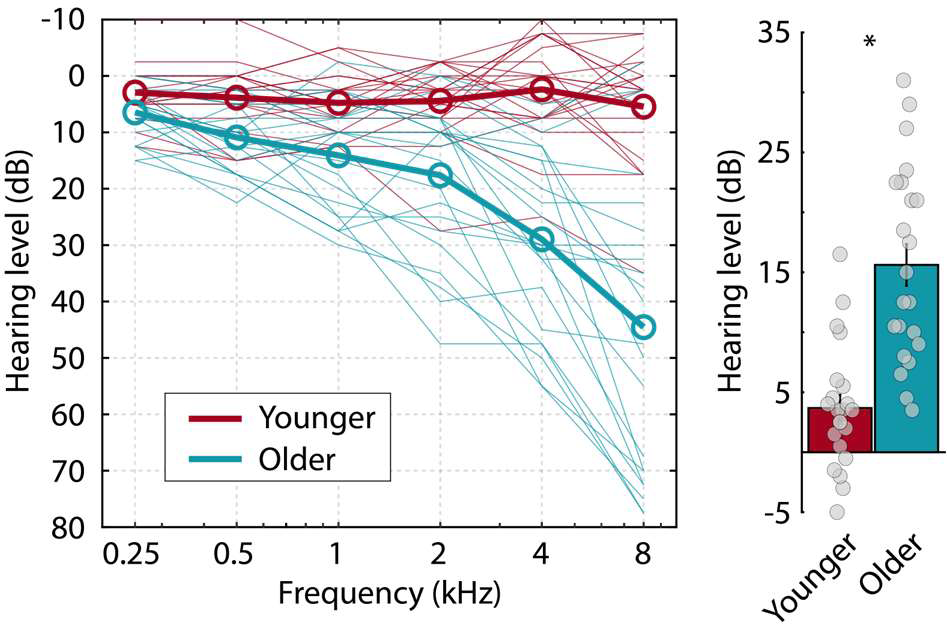
Pure-tone audiometric thresholds. Left: Pure-tone audiometric thresholds for all frequencies. The thin lines reflect data from each individual participant. Thick lines reflect the mean across participants. Right: Pure-tone average hearing threshold (mean across 0.25, 0.5, 1, 2, and 4 kHz). Dots reflect the pure-tone average threshold for individual participants. *p≤0.05.

### Experimental procedures for noise-burst stimulation

The current study is concerned with the reliability of the neural-tracking response during story listening. To obtain a “benchmark” against which to compare neural-tracking responses during speech listening, participants passively listened to 132 repetitions of a 0.1-s white-noise burst (0.01 s fade-in and 0.01 s fade-out) in a separate ∼3.5 min block of stimulation. The noise burst was presented with a mean onset-to-onset interval of 1.5 s (randomly selected between 1.2 and 1.8 s). We expected high reliability for responses to the noise burst. Noise-related reliability is expected to provide an upper reliability bound for story-related neural responses because noise bursts are spectrally broad, eliciting broad activity across auditory cortex.

### Experimental procedures for story listening

Across the two sessions, participants listened to six stories from the story-telling podcast The Moth (https://themoth.org/). The Moth podcast features spoken stories about human experiences and life events. The stories are intended to create an absorbing and enjoyable experience for the listener. The Moth stories mirror speech in everyday life and, unlike an audiobook, include disfluencies, filler-words, sentence fragments, corrections, unintentional pauses, and more flexible grammar (Tree, 1995; Bortfeld et al., 2001). In the current study, we used the following six stories, each of which was approximately 7 min in duration: *The Loudest Whisper* by Devan Sandiford, *Speech Writers Lament* by Karen Duffin, *Lego Crimes* by Micaela Blei, *Do The Dishes And Leave* by Mitch Donaberger, *Priceless Mangos* by Saya Shamdasani, and *Teacher Talent Show* by Tim Manley. Participants also listened to two moderately engaging and easily comprehensible story-book stories made for listeners at any level (Irsik et al., 2022a; Mathiesen et al., 2023). These stories were adapted from two print story books, *Wave* by D.M. Ouellet and *Alibi* by Kristin Butcher, aimed at reluctant readers. The stories are high interest but have a simple vocabulary and a linear plot. They are somewhat similar to audiobooks, which have been used frequently for neural speech tracking (Presacco et al., 2016b, a; Broderick et al., 2018; Lesenfants et al., 2019; Broderick et al., 2022). The two stories were recorded in-lab by a male native English speaker and each story was about 10 min in duration.

In each of the two sessions, participants listened to four The Moth stories and one Story Book story. Two of The Moth stories were presented under clear conditions (i.e., without background noise), whereas 12-talker babble at a signal-to-noise ratio (SNR) of 9 dB was added to the other two The Moth stories (Bilger, 1984; Bilger et al., 1984; Wilson et al., 2012). Speech in background babble at 9 dB SNR is still highly intelligible (∼90% of words; Irsik et al., 2022b), but may require more attention and effort by the listener than clear speech (Herrmann and Johnsrude, 2020; Yasmin et al., 2023). Moderate levels of background noise has also been shown to increase the neural tracking response (Yasmin et al., 2023). Story-book stories were presented under clear conditions. All stories were normalized to the same overall root-mean-square amplitude.

Stories were separated into three categories, henceforth referred to as SM stories (as in Same in both sessions, The Moth), DM stories (as in Different in both sessions, The Moth), and DB stories (as in Different in both sessions, Story Book; Figure 2A). SM stories were repeated in session 1 and session 2 to investigate test-retest reliability in a strict sense. We used The Moth stories by Devan Sandiford and Karen Duffin. SM stories were presented in the first two blocks of each session to reduce variance that may be associated with the session duration. The story by Devan Sandiford was presented under clear conditions both sessions, whereas the story by Karen Duffin was presented in background babble at 9 dB SNR in both sessions. The application of background babble was not counterbalanced across stories/sessions because this would have interfered with examining reliability. Story order was counter-balanced across the two sessions, such that if the story by Devan Sandiford was presented in the first block in session 1, it was presented in the second block of session 2, and vice versa (Figure 2A).

**Figure 2:**
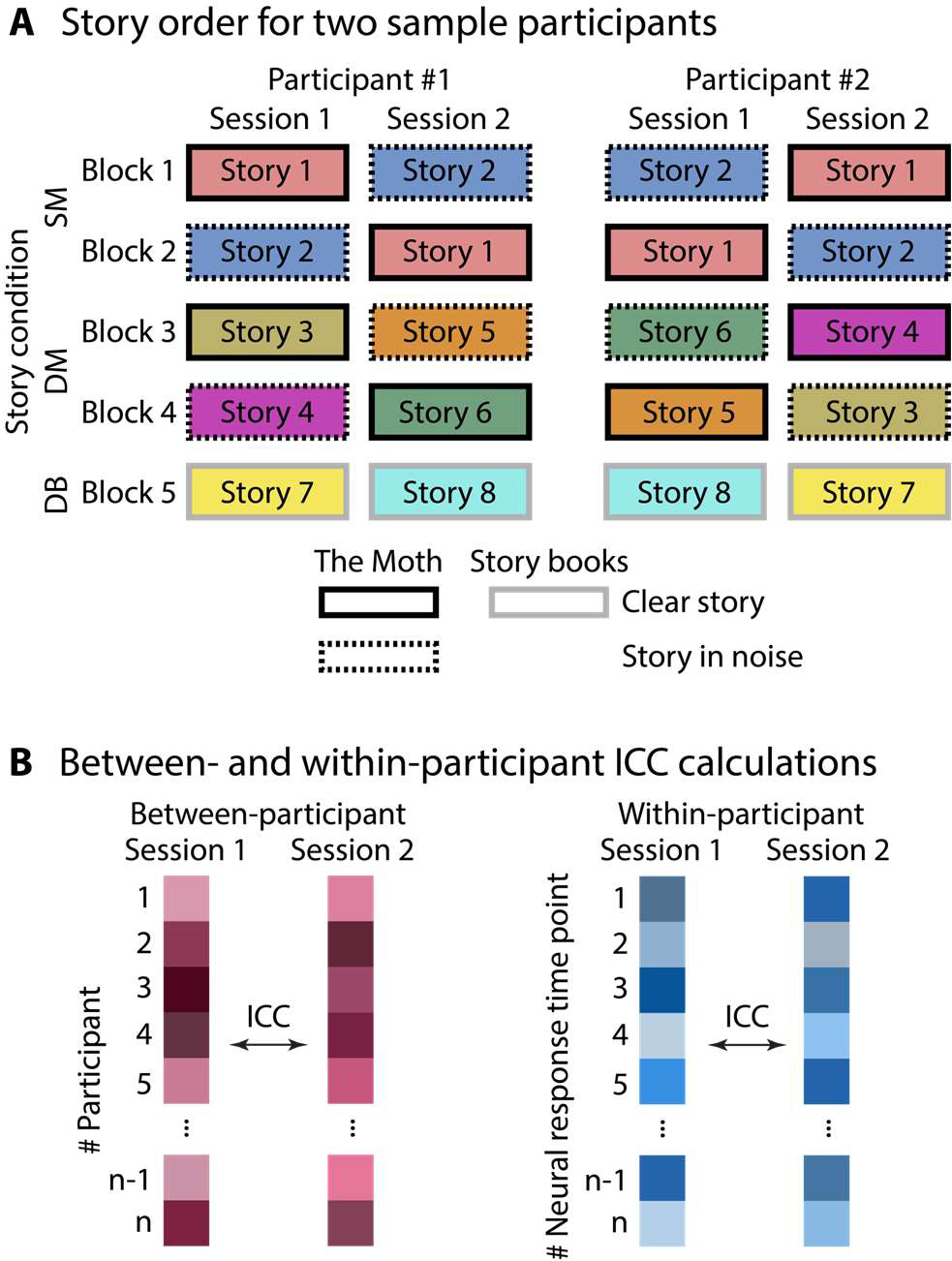
Experimental design and measures. **A:** Schematic of experimental design. Story order for two sample participants. Colors code the eight different stories and edge lines specify the speech-clarity conditions and story origin (solid – clear, dashed – babble noise; black – The Moth podcast, gray – Story Book). SM stories repeated across the two sessions (SM - same, The Moth), whereas DM stories did not repeat across sessions and were spoken by different speakers (DM – different, The Moth). SM and DM stories were sampled from The Moth podcast. DB stories mirror audiobook recordings (DM – story book) and were spoken by the same speaker. **B:** Schematic of the two types of intra-class correlation (ICC) calculations: between-participant ICC, and within-participant ICC. For between-participant ICC, observations are the participants, whereas, for within-participants ICC, observations are the neural response time points of a participant. Different colors schematically represent different response magnitudes.

DM stories were not repeated in session 1 and session 2. Two different stories from The Moth were presented in each session. DM stories enable the investigation of the generalizability of the neural-tracking response across sessions, stories, and speakers. In addition, by comparing the neural-tracking responses across stories within a session, we can investigate the generalizability across stories and speakers within a session. For DM stories, the stories by Micaela Blei, Mitch Donaberger, Saya Shamdasani, and Tim Manley were used. Two DM stories were always presented in blocks 3 and 4 of an experimental session (Figure 2A). One of the DM stories per session was presented under clear conditions, whereas the other DM story was presented with added background babble at a 9-dB SNR. Story order and speech-clarity conditions (clear, background noise) were counter-balanced across sessions and participants (Figure 2A). Counterbalancing enables us to make clearer inferences, compared to SM stories, about the effect of background babble on the neural-tracking response and its generalizability.

DB stories were story-book stories and mirrored more closely an audiobook than The Moth stories. One DB story was presented in each experimental session, always in block 5. The specific story-book story presented in a session was counter-balanced across participants (Figure 2A). DB stories enable us to investigate the generalizability of neural speech tracking across sessions, but reveal the effect of maintaining the same speaker. Moreover, the analysis of responses to DB stories provides insights into the reliability of neural tracking under more standardized conditions, where recordings are made in-lab by the same speaker. Such stories are more easily created in high numbers if needed for clinical assessments.

After each story, participants answered ten comprehension questions about the story. Each comprehension question comprised four response options (chance level = 25%). Participants also rated the degree to which they were absorbed by and enjoyed the story, using four and two items from previous work, respectively (Kuijpers et al., 2014; Herrmann and Johnsrude, 2020; absorption: “I felt absorbed by the story.”, “When I finished listening, I was surprised to see that time had gone by so fast.”, “I felt connected with the main character(s) of the story.”, “I could imagine what the world in which the story took place looked like.”; enjoyment: “I listened to the story with great interest.”, “I thought it was an exciting story.”). Participants further rated the degree to which they had to invest effort to comprehend the story using two items (“I had to invest effort to understand what was said.”, “Understanding the speaker was hard.”). Absorption, enjoyment, and effort were rated on a 7-point scale ranging from 1 (completely disagree) to 7 (completely agree).

### Analysis of behavioral story-listening data

Absorption, enjoyment, and effort rating scores were linearly re-scaled such that they range from 0 to 1 in order to facilitate interpretation and similarity to comprehension scores. For each participant, scores and ratings were separately averaged across story comprehension, effort, enjoyment, and absorption items.

Responses to DM stories were analyzed to assess the effects of Speech Clarity and Age Group on the four measures. To this end, a repeated measures analysis of variance (rmANOVA) was calculated, separately for story comprehension, effort, enjoyment, and absorption. Session (session 1, session 2) and Speech Clarity (clear, noise) were within-participants factors, whereas Age Group (younger, older) was a between-participants factor. Session was included to explicitly examine whether participants changed their judgement criteria across multiple lab visits, although we did not expect an effect of session due to counterbalancing story order and speech-clarity conditions (Figure 2A).

Responses to SM stories were analyzed to investigate the effects of Session and Age Group on the four measures. A rmANOVA was calculated separately for each behavioral measure (comprehension, effort, enjoyment, and absorption) using Session (session 1, session 2) as a within-participants factor and Age Group (younger, older) as a between-participants factor. Analyses were separately conducted for the clear story and the story in noise, because each speech-clarity condition for SM stories was associated with a specific story to facilitate reliability analyses as outlined above.

Responses to DB stories were analyzed to investigate the effects of Session and Age Group on the four measures for the audiobook-like stories. A rmANOVA was calculated separately for each behavioral measure (comprehension, effort, enjoyment, and absorption) using Session (session 1, session 2) as a within-participants factor and Age Group (younger, older) as a between-participants factor.

We also calculated the difference between enjoyment and absorption between The Moth stories (clear, session 1, SM & DM stories) and story-book stories (clear, session 1, DB stories) using a rmANOVA with Story Condition (The Moth, Story Book) as a within-participants factor and Age Group (younger, older) as a between-participants factor.

### Electroencephalography recordings and preprocessing

Electroencephalographical signals were recorded from 16 scalp electrodes (Ag/Ag–Cl-electrodes; 10-20 placement) and the left and right mastoids using a BioSemi system (Amsterdam, The Netherlands). The sampling frequency was 1024 Hz with an online low-pass filter of 208 Hz.

Electrodes were referenced online to a monopolar reference feedback loop connecting a driven passive sensor and a common-mode-sense (CMS) active sensor, both located posteriorly on the scalp.

Offline analysis was conducted using MATLAB software. An elliptic filter was used to suppress power at the 60-Hz line frequency. Data were re-referenced by averaging the signal from the left and right mastoids and subtracting the average separately from each of the 16 channels. Data were filtered with a 0.7-Hz high-pass filter (length: 2449 samples, Hann window) and a 22-Hz low-pass filter (length: 211 samples, Kaiser window).

For the one block during which noise bursts were presented, EEG data were divided into epochs ranging from −1 to 2 s time-locked to noise onset and down-sampled to 512 Hz. Independent components analysis (runica method, Makeig et al., 1996; logistic infomax algorithm, Bell and Sejnowski, 1995; Fieldtrip implementation Oostenveld et al., 2011) was used to identify and remove components related to blinks and horizontal eye movements. Epochs for which the signal range exceeded 100 µV in any of the EEG electrodes were excluded from analysis.

For the five blocks during which stories were presented, EEG data were segmented into time series time-locked to story onset and down-sampled to 512 Hz. Independent components analysis was used to remove signal components reflecting blinks and eye movement (Bell and Sejnowski, 1995; Makeig et al., 1996 Oostenveld et al., 2011). Additional artifacts were removed after the independent components analysis by setting the voltage for segments in which the EEG amplitude varied more than 80 µV within a 0.2-s period in any channel to 0 µV (cf. Dmochowski et al., 2012; Dmochowski et al., 2014; Cohen and Parra, 2016; Irsik et al., 2022b; Yasmin et al., 2023). Data were low-pass filtered at 10 Hz (251 points, Kaiser window) because neural signals in the low-frequency range are most sensitive to acoustic features (Di Liberto et al., 2015; Zuk et al., 2021; Yasmin et al., 2023).

For one younger participant, EEG recordings for one SM story and, for another younger participant, EEG recordings for one DM story could not be analyzed because of a technical error during recording. EEG data from these participants were excluded for analyses that required the availability of data from both sessions but were otherwise included.

### Analysis of event-related potentials to noise bursts

Time courses were averaged across trials, focusing on −0.15 to 0.5 s epochs time-locked to noise onset. Response time courses were baseline-corrected by subtracting the mean amplitude within the −0.15 to 0 s time window from the amplitude at each time point. Neural responses were also averaged across a fronto-central electrode cluster (F3, Fz, F4, C3, Cz, C4) known to be sensitive to neural activity originating from auditory cortex (Näätänen and Picton, 1987; Picton et al., 2003; Herrmann et al., 2018; Irsik et al., 2021).

Data analysis focused on the amplitude in the 0.03–0.06 s and the 0.08–0.12 s time windows that correspond to the P1 and N1 components of the event-related potentials (Picton et al., 1974; Näätänen and Picton, 1987; Herrmann et al., 2016; Herrmann et al., 2018). Mean amplitudes within a time window were used as dependent measure in a rmANOVA. Predictors were the within-participants factor Session (session 1, session 2) and the between-participants factor Age Group (younger, older).

Reliability of neural responses was calculated in two ways, focusing on between-participant reliability and within-participant reliability. For between-participant reliability (Figure 2B, left), intra-class correlation (ICC) was calculated as the two-way mixed effects, absolute agreement, single rater (ICC(2,1); Shrout and Fleiss, 1979; McGraw and Wong, 1996; Koo and Li, 2016). Specifically, amplitudes were averaged and ICC was calculated for 0.5 s sliding windows centered on each time point, separately for each age group. That is, the mean time-window response in session 1 and session 2 were the measures, while participants were the observations in this analysis. This resulted in one ICC time course per age group. ICC values indicate excellent, good, moderate, or poor reliability if they are greater than 0.9, between 0.75 and 0.9, between 0.5 and 0.75, or below 0.5, respectively (Koo and Li, 2016). Note that between-participant ICC provides a descriptive measure for which no statistical test is available because ICC values are not calculated for each participant, but across participants.

For within-participant reliability (Figure 2B, right), we calculated ICC values (ICC(2,1); Shrout and Fleiss, 1979; Koo and Li, 2016), separately for each participant as the reliability of the time course ranging from 0 to 0.4 s across sessions. That is, response time courses in session 1 and session 2 of the same participant were the two measures, while individual time points were the observations in this analysis. The 0-0.4 s time window was chosen because it comprises the major deflections of the event-related potential (Figure 3). This resulted in one ICC value for each participant. Age-group differences in within-participant ICC were assessed using an independent-samples t-test.

**Figure 3:**
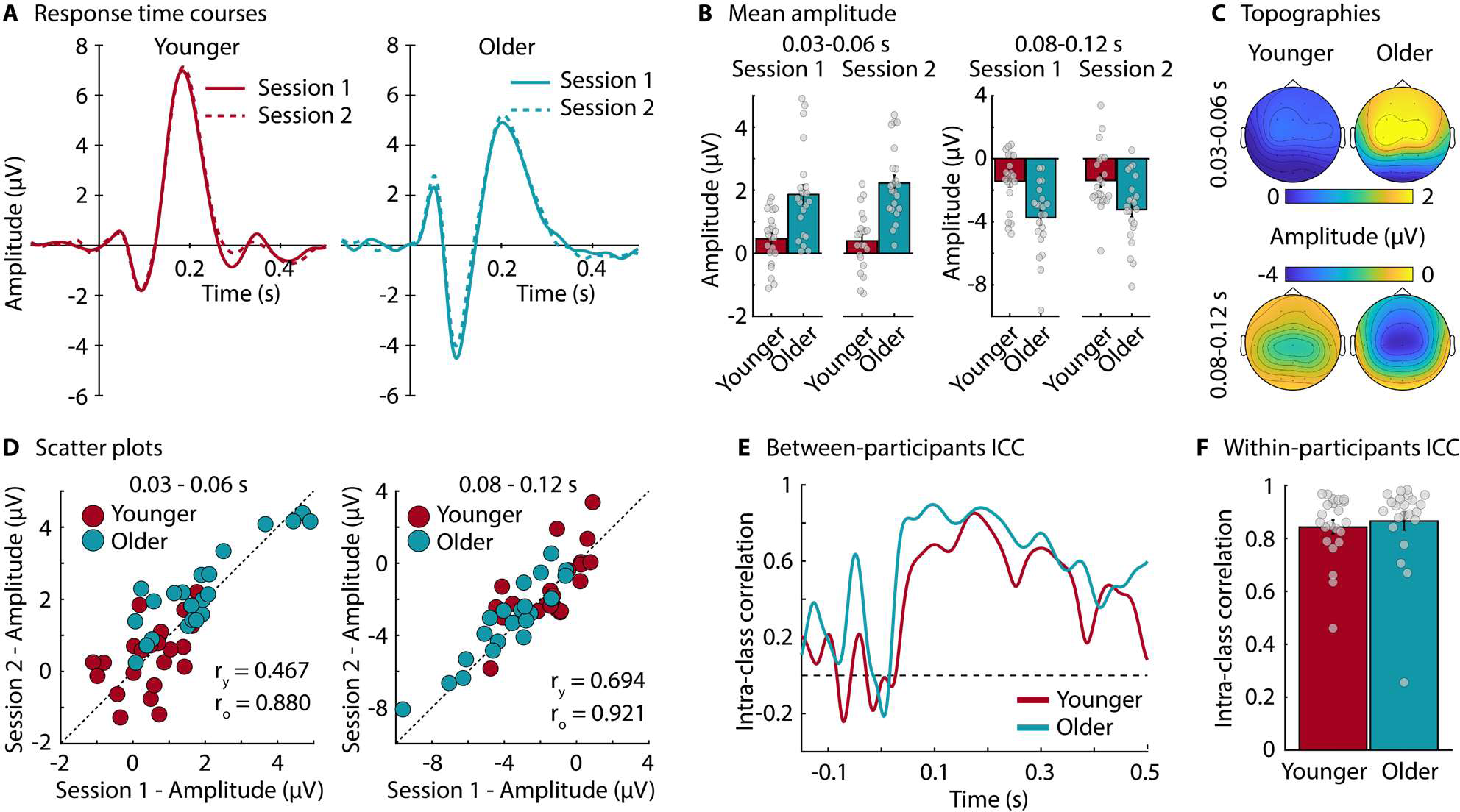
Neural responses and reliability for noise bursts. **A:** Time courses of neural responses to noise bursts and topographical distributions for the 0.03–0.06 s and the 0.08–0.12 s time windows (averaged across sessions). **B:** Mean neural responses for the 0.03–0.06 s and the 0.08–0.12 s time windows. Dots reflect data from individual participants. **C:** Topographical distributions (averaged across sessions). **D:** 45-degree scatter plots for neural responses in the 0.03–0.06 s and the 0.08–0.12 s time windows. Pearson correlations are provided within each plot. The subscripts y and o indicate correlations for younger and older adults, respectively. **E:** Between-participant ICC time courses for younger and older adults. ICC was calculated for 0.05 s time windows centered on each time point. **F:** Within-participant ICC, considering the time course from 0 s to 0.4 s. Dots reflect data from individual participants.

### Calculation of neural tracking response and EEG reconstruction accuracy during story listening

We used a forward model based on the linear temporal response function (TRF; Crosse et al., 2016; Crosse et al., 2021) to quantify the relationship between the amplitude envelope of a story and EEG activity. To this end, a cochleogram was calculated for each story using a simple auditory-periphery model with 30 auditory filters (McDermott and Simoncelli, 2011). The resulting amplitude envelope for each auditory filter was compressed by 0.6 to simulate inner ear compression (McDermott and Simoncelli, 2011). Such a computationally simple peripheral model has been shown to be sufficient, as compared to complex, more realistic models, for envelope-tracking approaches (Biesmans et al., 2017). Amplitude envelopes were averaged across auditory filters and low-pass filtered at 40-Hz filter (Butterworth filter). To obtain the amplitude-onset envelope, we calculated the first derivative and set all negative values to zero (Hertrich et al., 2012; Fiedler et al., 2017; Fiedler et al., 2019; Yasmin et al., 2023). The amplitude-onset envelope was down-sampled to match the sampling of the EEG data.

For the analysis, 100 30-s data snippets (Crosse et al., 2016; Crosse et al., 2021) were extracted randomly from the EEG data and corresponding amplitude-onset envelope per story and session. Each of the 100 EEG and envelope snippets were held out once as a test dataset, while the remaining non-overlapping EEG and envelope snippets were used as training datasets. That is, for each training dataset, linear regression with ridge regularization was used to map the amplitude-onset envelope onto the EEG activity to obtain a TRF model for lags ranging from 0 to 0.4 s (Hoerl and Kennard, 1970; Crosse et al., 2016; Crosse et al., 2021). The ridge regularization parameter lambda (λ), which prevents overfitting, was set to 10 based on previous work (Fiedler et al., 2017; Fiedler et al., 2019; Yasmin et al., 2023). Note that cross-validation to estimate λ yielded qualitative similar results compared to a λ of 10. Pre-selection of λ based on previous work avoids extremely low and high λ on some cross-validation iterations and avoids substantially longer computational time that may be unfeasible in clinical contexts. Pre-selection of λ may also be required clinically because assessments times need to be short, which limit the amount of data recorded (Crosse et al., 2021). The TRF model calculated for the training data was then used to predict the EEG signal for the held-out test dataset. The Pearson correlation between the predicted and the observed EEG data of the test dataset was used as a measure of EEG reconstruction accuracy (Crosse et al., 2016; Crosse et al., 2021). Model estimation and reconstruction accuracy were calculated separately for each of the 100 data snippets per story and session, and reconstruction accuracies were averaged across the 100 snippets. To investigate the neural-tracking response directly in addition to the reconstruction accuracy, we also calculated TRFs for each training dataset for a broader set of lags, ranging from −0.15 to 0.5 s, to enable similar analyses as for traditional event-related potentials (Yasmin et al., 2023). TRFs were averaged across the 100 training datasets.

Data analyses of response magnitude differences and reliability/generalizability of neural responses focused on a fronto-central electrode cluster (F3, Fz, F4, C3, Cz, C4) known to be most sensitive to auditory cortex activity (Näätänen and Picton, 1987; Picton et al., 2003; Irsik et al., 2021). TRFs and reconstruction accuracies were averaged across the electrodes of this fronto-central electrode cluster prior to further analysis.

### Analysis of the effects of speech clarity and age group on neural response magnitude

Before investigations into the reliability of neural responses, we first aimed to examine differences in response amplitude related to speech clarity and age group. To this end, we focused analyses on TRFs and reconstruction accuracy for DM Stories, for which speech-clarity conditions were counter-balanced across stories and sessions. TRFs and reconstruction accuracies were averaged across stories and sessions, separately for clear stories and stories in babble noise. Analyses of the TRF focused on P1 and N1 amplitudes as the average in the 0.03–0.06 s and the 0.9–0.13 s time windows, respectively. A slightly later time window was chosen for the N1 in response to speech compared to the N1 response to the noise bursts, because our and previous data indicate later N1 latencies for speech (Hertrich et al., 2012; Fiedler et al., 2019; Yasmin et al., 2023; possibly due to more graded acoustic onsets). A rmANOVA was calculated separately for TRF P1, TRF N1, and reconstruction accuracy, using the within-participants factor Speech Clarity (clear, noise) and the between-participants factor Age Group (younger, older).

### Analysis of reliability of neural responses during story listening

Analyses of the reliability of the neural responses during story listening focused on SM stories, for which the identical stories (one clear, one with added babble) were presented in both sessions. Reliability analyses were calculated in two ways, similar to the neural responses to noise bursts, focusing on between-participant reliability and within-participant reliability (Figure 2B).

For between-participant reliability, ICC was calculated as the two-way, mixed effects, absolute agreement (ICC(2,1); Shrout and Fleiss, 1979; Koo and Li, 2016). TRF amplitudes were averaged and ICC was calculated for 0.5 s sliding windows centered on each time point, separately for the clear story and the story with added background babble, and separately for the two age groups. The mean time-window response in session 1 and session 2 were the measures in this ICC analysis, while participants were the observations (Figure 2B, left). This resulted in one ICC time course per speech-clarity condition and age group (statistical analyses cannot be conducted for between-participant ICC).

For within-participant reliability (Figure 2B, right), the TRF time courses (ranging from 0 to 0.4 s) of the two sessions were used to calculate ICC, separately for each participant and speech-clarity condition. Response time courses in the two sessions were the measures in this ICC analysis, while individual time points were the observations. Age-group differences in within-participant reliability (ICC) were assessed using an independent-samples t-test, separately for the clear story and the story in background babble (speech-clarity conditions were not directly compared because we did not counter-balance speech clarity across stories to reduce unwanted variance).

We also calculated between-participant reliability using ICC for reconstruction accuracy, separately for both speech-clarity conditions and both age groups. For this ICC analysis, the correlation values calculated using the leave-one-out procedure for session 1 and session 2, described above, were the measures, and participants were the observations (Figure 2B, left). This resulted in one ICC value per speech-clarity condition and age group. Again, statistical analyses cannot be conducted for between-participant ICC.

### Generalizability of neural-speech tracking responses

Generalizability of story-related neural responses was investigated within sessions and between sessions. For the investigation of generalizability across stories and speakers within a session, we focused only on session 1 to avoid influences of the repetition of SM stories. ICC was calculated separately for clear stories and stories with background babble (i.e., using one SM and one DM story each), and separately for the two age groups. To this end, TRF amplitudes were averaged and between-participant ICC was calculated for 0.5-s sliding windows centered on each time point. Between-participant ICC was also calculated for reconstruction accuracy, separately for each speech-clarity condition and age group. We further calculated within-participant ICC, for which the TRF time courses (ranging from 0 to 0.4 s) of the two stories of the same speech-clarity condition (clear, noise) were used to calculate ICC, separately for each participant. A rmANOVA was calculated using the within-participant ICC as the dependent variable, and Speech Clarity (clear, noise) and Age Group (younger, older) as independent variables.

For the investigation of the generalizability across sessions, stories, and speakers, ICC of neural responses to DM stories was calculated similarly as the reliability calculation for neural responses to SM stories. That is, TRF amplitudes were averaged and ICC was calculated for 0.5-s sliding windows centered on each time point, separately for clear stories and stories with added background babble, and separately for the two age groups. Between-participant ICC was also calculated for reconstruction accuracy, separately for both speech-clarity conditions and both age groups. We further calculated within-participant ICC, for which the TRF time courses (ranging from 0 to 0.4 s) of the responses to stories in the two sessions were used to calculate ICC separately for each participant and speech-clarity condition. A rmANOVA was calculated using the within-participant ICC as the dependent variable, and Speech Clarity (clear, noise) and Age Group (younger, older) as independent variables.

For the investigation of the generalizability of the neural-tracking response across sessions and speakers for audiobook-like stories, DB stories were used. Between-participant ICC time courses were calculated by averaging TRF amplitudes and computing ICC for 0.5-s sliding windows centered on each time point, separately for the two age groups. Between-participant ICC was also calculated for reconstruction accuracy, separately for both age groups. Within-participant ICC was further calculated (using the TRF time courses from 0 to 0.4 s) for each participant. An independent samples t-test was used to compare within-participant ICC between age groups.

### Comparisons among reliability and generalizability

To compare whether ICC differed between different assessment types, we calculated a rmANOVA using within-participant ICC values as the dependent measure and Assessment Type as within-participants factor with five levels: ERP reliability, TRF reliability (SM stories across sessions), TRF generalizability within a session (SM stories vs DM stories), TRF generalizability across sessions (DM stories), and TRF generalizability across session for stories spoken by the same person (DB stories). Age Group (younger, older) was included as a between-participants factor. For this analysis, we focused on clear stories because the noise bursts for the ERPs were also presented under clear conditions. Between-participant ICC could not be assessed statistically, because it is calculated across participants and no participant-specific ICC data points are available.

### Statistical analyses

All data analyses described were carried out using MATLAB (MathWorks) and JASP software (JASP, 2022; version 0.16.4.0). Effect sizes for rmANOVAs and t-tests are reported using omega squared (ω^2^) and Cohen’s d (d), respectively. Significant effects or interactions in rmANOVAs were resolved using post hoc tests with Holm’s method for multi-comparison corrections (Holm, 1979).

## Results

### Neural responses to noise bursts

Responses to noise bursts in the 0.03–0.06 s and in the 0.08–0.12 s time windows were larger for older compared to younger adults (effect of Age Group: 0.03–0.06 s: F_1,42_ = 26.616, p = 6.3 · 10^-6^, ω^2^ = 0.230; 0.08–0.12 s: F_1,42_ = 12.592, p = 9.7 · 10^-4^, ω^2^ = 0.119; Figure 2), but there was no difference between sessions (effect of Session: ps > 0.1) and no Session × Age Group interaction (ps > 0.05).

Between-participant reliability of the neural responses was moderate to good for younger adults (ICC ∼0.6-0.8) and good for older adults (ICC >0.8) in the 0.05–0.25 s time window (Koo and Li, 2016; Figure 3E). Mean within-participant reliability, considering the time course from 0 s to 0.4 s, was good for both younger (ICC = 0.84) and older adults (ICC = 0.87), and did not differ between age groups (t_42_ = 0.526, p = 0.601, d = 0.159; Figure 3F).

### Behavioral data for story listening

We first focused on DM stories, that is, those that were not repeated and for which speech-clarity conditions were counterbalanced across sessions and stories. The behavioral data for DM stories enabled us to investigate the effects of Speech Clarity and Age Group on story comprehension, listening effort, enjoyment, and absorption as well as whether participants changed how they rated the metrics from session 1 to session 2 (Figure 4A). Listening effort was higher for stories in background noise compared to clear speech (effect of Speech Clarity: F_1,42_ = 46.933, p = 2.4 · 10^-8^, ω^2^ = 0.249). No other effects nor interactions, for any of the measures, were significant (ps > 0.05). Hence, enjoyment, absorption, and comprehension were not significantly affected by background noise (Herrmann and Johnsrude, 2020), there were no evident differences between age groups (Mathiesen et al., 2023), and there was no overall tendency to rate the measures differently in the second compared to the first session (no effects of Session).

**Figure 4:**
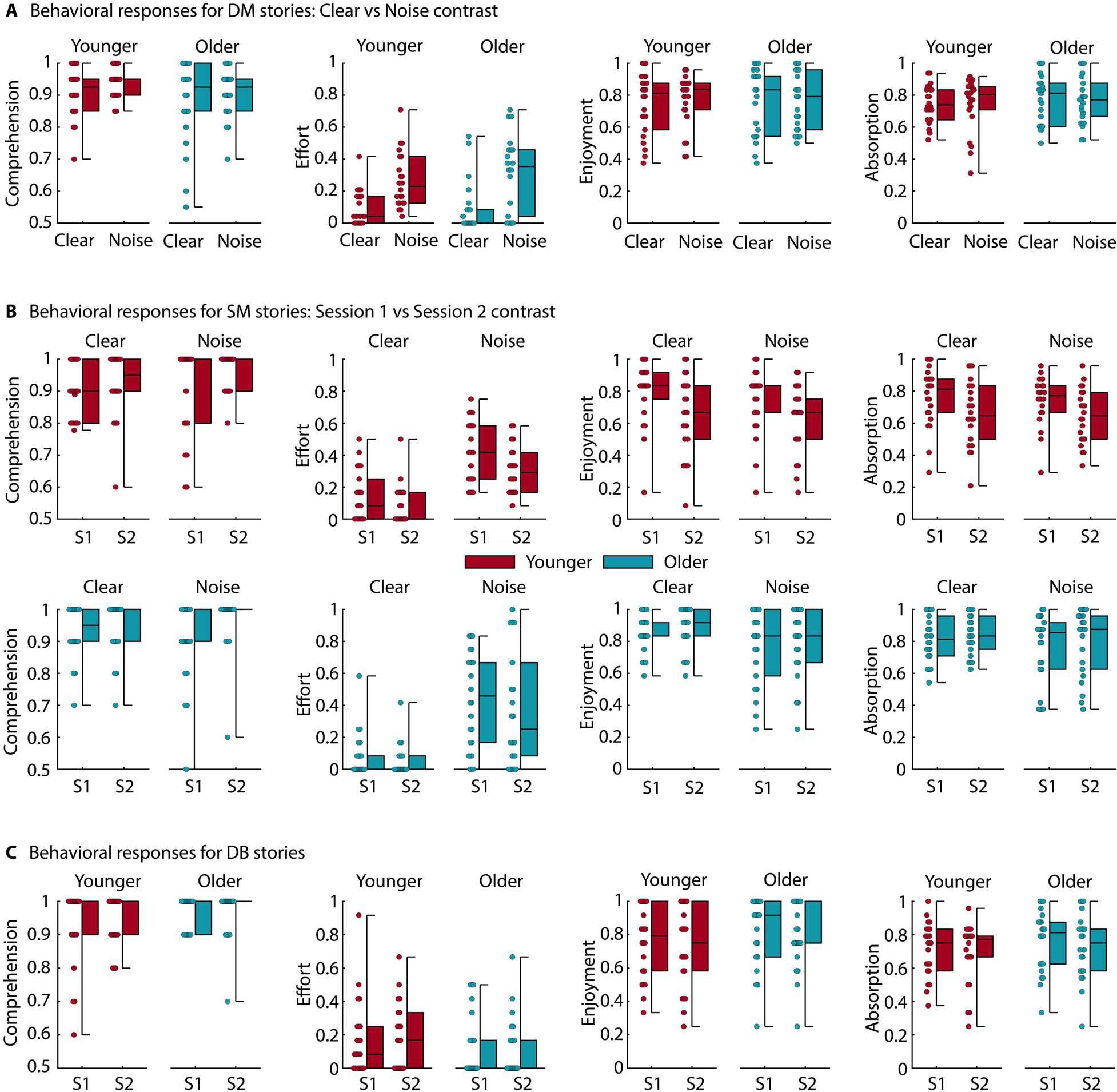
Behavioral data for story listening. **A:** Story comprehension, listening effort, enjoyment, and absorption scores for DM stories (different The Moth stories in the two sessions), which enable comparisons between clear stories and stories in babble. **B:** Story comprehension, listening effort, enjoyment, and absorption scores for SM stories (same The Moth stories in both sessions), which enable comparison between sessions. First row in panel B shows data from younger adults, whereas the second row shows the data for older adults. **C:** Story comprehension, listening effort, enjoyment, and absorption scores for DB stories (different story-book stories in the two sessions). Box plots are shown as well as data point from individual participants. S1 – session 1, S2 – session 2.

Analyses for SM stories (i.e., identical stories in both sessions) aimed to investigate the effects of Session and Age Group on story comprehension, listening effort, enjoyment, and absorption, separately for the clear story and the story in babble (Figure 4B). Story comprehension did not differ between sessions under clear conditions (p > 0.2), but comprehension was higher in session 2 than session 1 under babble conditions (F_1,42_ = 11.618, p = 0.001, ω^2^ = 0.089). There were no effects of Age Group or interactions for comprehension (ps > 0.2). Effort decreased from session 1 to session 2 for the clear story (F_1,42_ = 5.029, p = 0.030, ω^2^ = 0.022) and the story in noise (F_1,42_ = 4.689, p = 0.036, ω^2^ = 0.026). There were again no effects of Age Group or interactions (ps > 0.1). For enjoyment, the rmANOVA revealed a Session × Age Group interaction for the clear story (F_1,42_ = 14.948, p = 3.8 · 10^-4^, ω^2^ = 0.058) and the story in noise (F_1,42_ = 7.701, p = 0.008, ω^2^ = 0.022), such that enjoyment was lower in session 2 than session 1 for younger adults (clear: t_42_ = 4.353, p_Holm_ = 4.2 · 10^-4^, d = 0.806; noise: t_42_ = 2.970, p_Holm_ = 0.029, d = 0.486), but not for older adults (for both p_Holm_ > 0.5). Results for absorption mirrored those for enjoyment. A Session × Age Group interaction was observed for the clear story (F_1,42_ = 7.955, p = 0.007, ω^2^ = 0.029) and the story in noise (F_1,42_ = 9.050, p = 0.004, ω^2^ = 0.030), showing lower absorption for session 2 than 1 for younger (clear: t_42_ = 3.229, p_Holm_ = 0.012, d = 0.587; noise: t_42_ = 3.031, p_Holm_ = 0.025, d = 0.525), but not for older adults (for both p_Holm_ > 0.5). Enjoyment and absorption were also higher for older compared to younger adults for the clear story (effect of Age Group: enjoyment: F_1,42_ = 9.046, p = 0.004, ω^2^ = 0.086; absorption: F_1,42_ = 6.739, p = 0.013, ω^2^ = 0.063), but not the story in noise (ps > 0.05).

For DB stories, there were no differences between sessions nor between age groups for any of the measures (ps > 0.05). Moreover, story-book stories (DB stories) appeared to be as enjoyable and absorbing as The Moth stories (SM & DM stories; ps > 0.1; Figure 4C).

In sum, the behavioral data show that speech in babble noise increases listening effort, but comprehension is higher and listening effort reduced when individuals listen to the same story again a week or more later than when listening to it for the first time. Enjoyment and absorption also decreased with story repetition, but this was only the case for younger adults. Older adults appeared to similarly enjoy and be absorbed by the stories in both sessions.

### Age- and noise-related increases in neural-tracking response during story listening

Prior to analyses of neural-tracking reliability, we investigated the degree to which age group and speech clarity affect the TRF amplitude and reconstruction accuracy (Figure 5). These analyses focused on DM stories for which speech-clarity conditions were counter-balanced across stories and sessions.

**Figure 5:**
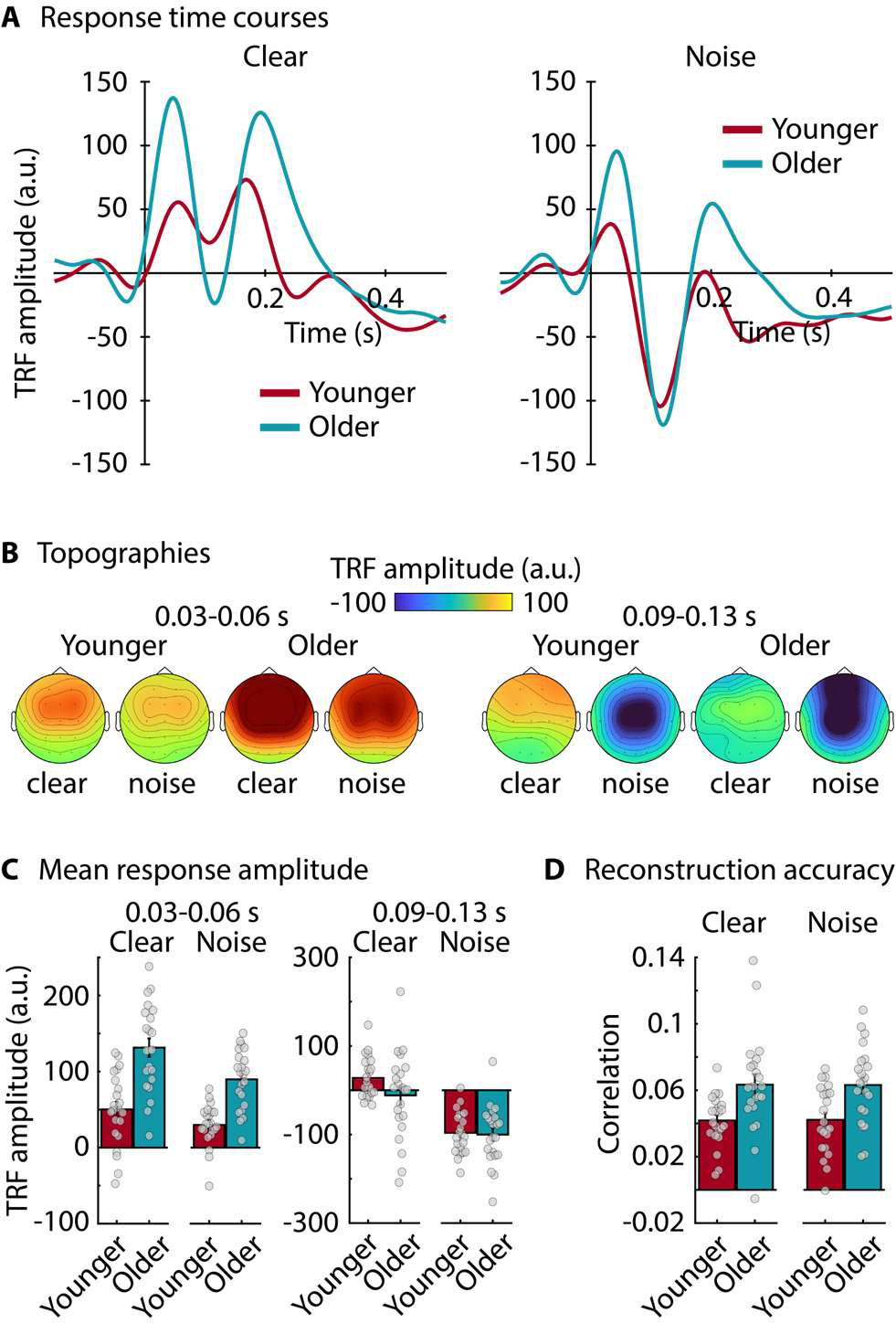
Effects of Speech Clarity and Age Group on TRF amplitude and EEG reconstruction accuracy using DM stories. **A:** Temporal response functions (TRF) for each speech-clarity condition (clear, noise) and age group (younger, older). **B:** Topographies for the mean TRF amplitude in the 0.03–0.06 s and the 0.09–0.13 s time windows. **C:** Bar graphs show the mean TRF amplitude in the 0.03–0.06 s and the 0.09–0.13 s time windows. Dots reflect data from individual participants. **D:** EEG reconstruction accuracy for each speech-clarity condition (clear, noise) and age group (younger, older). Dots reflect data from individual participants.

For the TRF analysis (averaged across stories and sessions) in the 0.03–0.06 s time window, amplitudes were larger in older compared to younger adults (effect of Age Group: F_1,42_ = 43.841, p = 5.1 · 10^-8^, ω^2^ = 0.333) and smaller for stories in background noise compared to clear stories (effect of Speech Clarity; F_1,42_ = 15.221, p = 3.4 · 10^-4^, ω^2^ = 0.106). The Speech Clarity × Age Group interaction was not significant (F_1,42_ = 1.786, p = 0.189, ω^2^ = 0.006; Figure 5C).

For the 0.09–0.13 s time window, TRF amplitudes were larger (i.e., more negative) for stories in babble compared to clear stories (effect of Speech Clarity: F_1,42_ = 104.036, p = 6.2 · 10^-13^, ω^2^ = 0.381; Figure 5A). There was no effect of Age Group (F_1,42_ = 1.498, p = 0.228, ω^2^ = 0.006) nor a Speech Clarity × Age Group interaction (F_1,42_ = 2.888, p = 0.097, ω^2^ = 0.011; Figure 5C).

The rmANOVA for EEG reconstruction accuracy revealed larger accuracies for older compared to younger adults (F_1,42_ = 14.027, p = 5.4 · 10^-4^, ω^2^ = 0.132). The effect of Speech Clarity (F_1,42_ < 0.001, p = 0.982, ω^2^ < 0.001) and the Speech Clarity × Age Group interaction (F_1,42_ = 0.005, p = 0.942, ω^2^ < 0.001) were not significant.

### Moderate reliability of neural-tracking response during story listening

The analyses reported in this section focus on SM stories and explore reliability in a strict sense (i.e., identical stories presented in both sessions). TRF amplitudes in the 0.03–0.06 s time window were larger for older compared to younger adults (clear: F_1,42_ = 31.290, p = 1.6 · 10^-6^, ω^2^ = 0.265; noise: F_1,42_ = 22.764, p = 2.3 · 10^-5^, ω^2^ = 0.206; Figure 6), mirroring the results for DM stories (Figure 5). There were no effects of Session, nor Session × Age Group interactions, nor any effects for the 0.09–0.13 s time window (ps > 0.15). EEG reconstruction accuracy was also greater in older compared to younger adults (clear: F_1,42_ = 15.826, p = 2.8 · 10^-4^, ω^2^ = 0.150; noise: F_1,42_ = 11.250, p = 0.002, ω^2^ = 0.109; Figure 6E), but there were no effects of Session nor Session × Age Group interactions (ps > 0.4).

**Figure 6:**
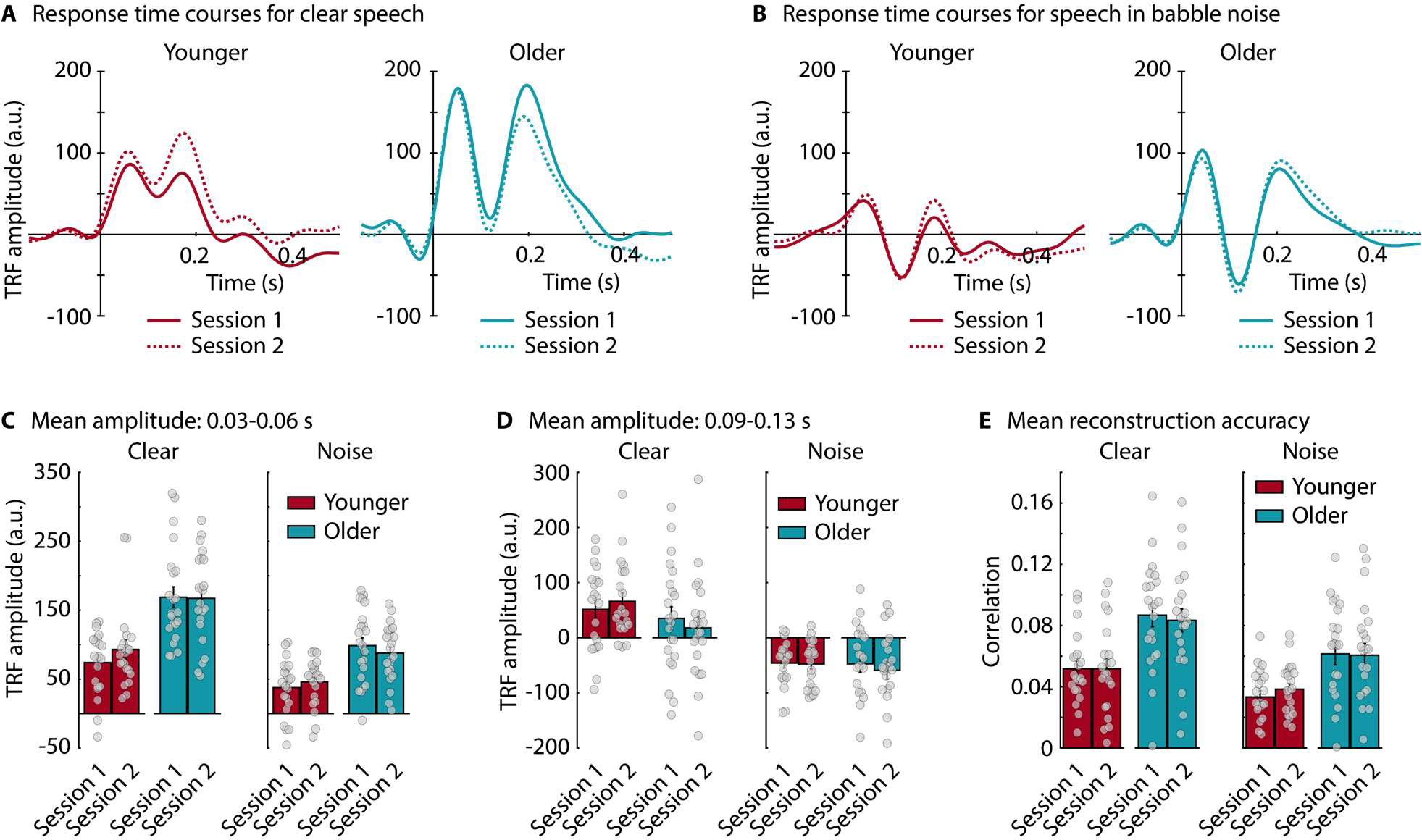
Neural responses in session 1 and 2 for SM stories. **A:** Temporal response functions (TRFs) for sessions and age groups for the clear story. **B:** TRFs for sessions and age groups for the story in babble. **C:** Mean TRF amplitude in the 0.03-0.06 s time window. Bars reflect the mean across participants and gray dot data from individual participants. Error bars reflect the standard error of the mean. **D:** Same as in panel C for the TRF amplitude in the 0.09-0.13 s time window. **E:** Same as in panel C for EEG reconstruction accuracy.

Figure 7 shows reliability data for the neural responses elicited by SM stories. Time courses in Figure 7A and B (middle) show the between-participant ICC for TRF amplitudes using 0.05-s sliding time windows. Peak ICC at around 0.1 to 0.2 s was about 0.72 for younger and older adults for the clear story, but below 0.6 for most other time points. Peak ICC for the story in babble noise was about 0.75 for older and 0.5 for younger adults. ICC values between 0.5 and 0.75 are indicative of moderate reliability (Koo and Li, 2016).

**Figure 7:**
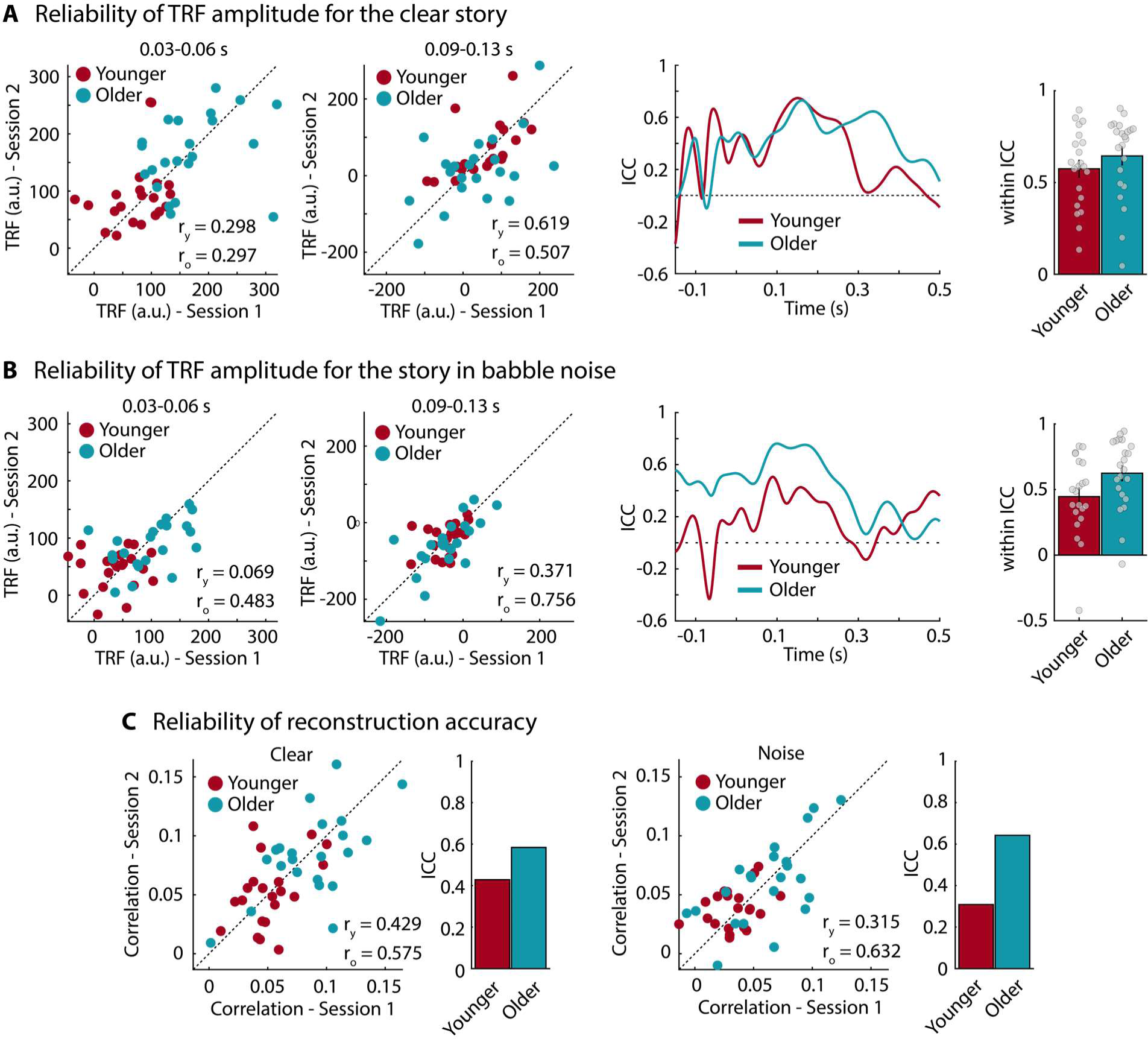
Reliability of neural responses for SM stories. **A:** Reliability for TRF amplitude for the clear story. Left: 45-degree scatter plots for TRF amplitude in the 0.03-0.06 s and the 0.09–0.13 s time windows. Pearson correlations are provided. The subscripts y and o indicate correlations for younger and older adults, respectively. Middle: Between-participant ICC time courses for both age groups (younger, older). ICC was calculated for 0.05 s time windows centered on each time point. Right: Within-participant ICC, considering the time course from 0-0.4 s. Bars reflect the mean and dots reflect data from individual participants. **B:** Same as in panel A for the story in babble noise. **C:** Reliability for EEG reconstruction accuracy for the clear story (left) and the story in noise (right). 45-degree scatter plots and between-participant ICC are shown.

Within-participant ICC – that is, the agreement of the 0-0.4-s time courses between the two sessions – was 0.58 (younger) and 0.65 (older) for the clear story and 0.45 (younger) and 0.62 (older) for the story in noise. Within-participant ICC was greater for older compared to younger adults for the story in noise (t_41_ = 2.058, p = 0.046, d = 0.628), but there was no age-group difference for the clear story (t_41_ = 1.051, p = 0.299, d = 0.321; Figure 7A,B right).

Between-participant ICC for reconstruction accuracy of the clear story was 0.43 for younger and 0.58 for older adults. For the story in babble noise, between-participant ICC for reconstruction accuracy was 0.3 for younger and 0.64 for older adults (Figure 7C).

### Across-story generalizability: Within-sessions focus

Generalizability of neural responses across stories within session 1 was investigated by calculating ICC across SM stories and DM stories. Between-participant ICC for the TRF amplitude was about 0.6 at ∼0.1 s where it peaked for clear stories and stories in noise (moderate; Koo and Li, 2016), although ICC for stories in noise for the younger adults was lower (<0.5; Figure 8A, left).

**Figure 8:**
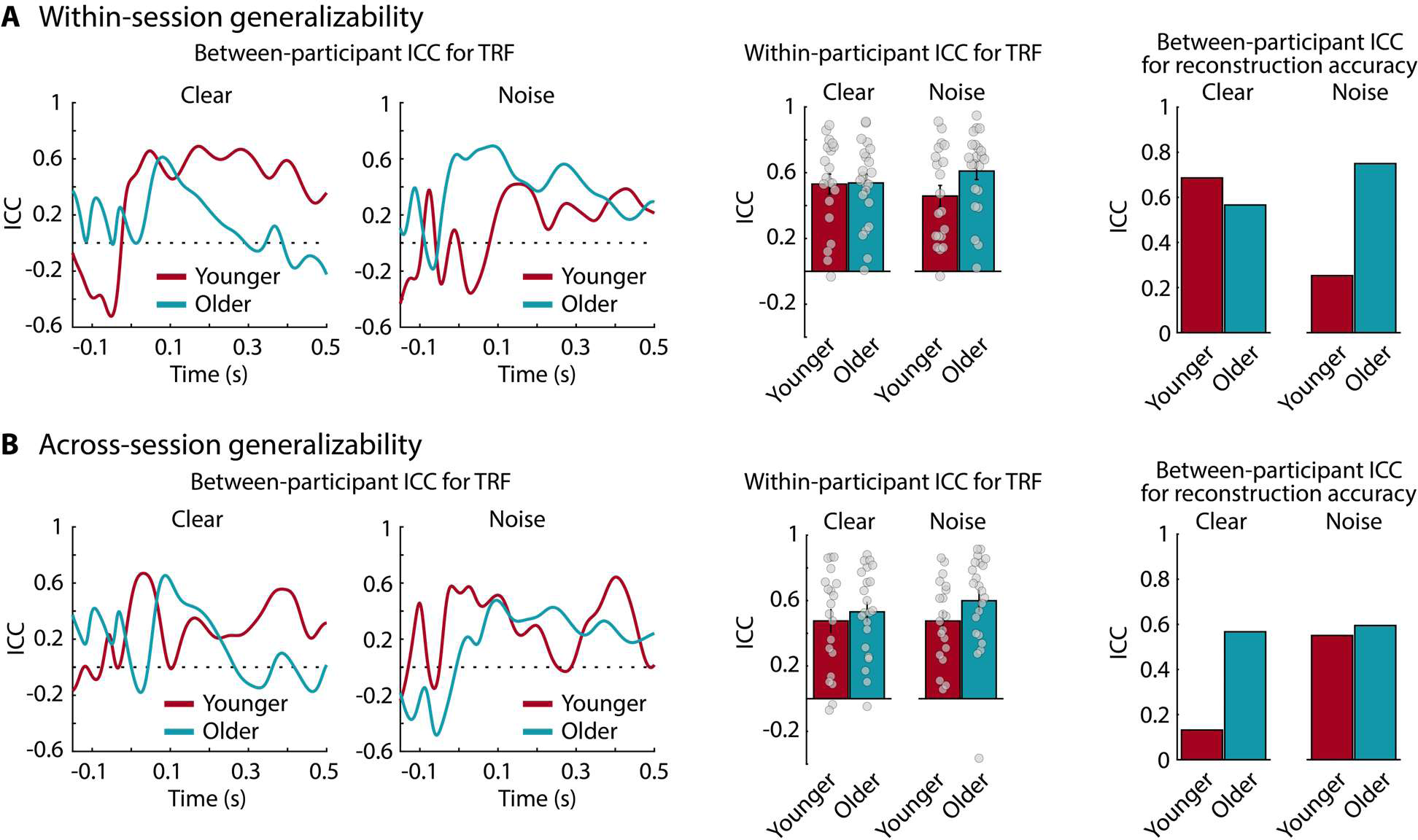
Generalizability of neural responses across stories, within and across sessions. **A:** Generalizability within a session (i.e., SM stories vs DM stories in session 1): Left: Between-participant ICC time courses for both speech-clarity conditions (clear, noise) and age groups (younger, older). ICC was calculated for 0.05 s sliding time windows centered on each time point. Middle: Within-participant ICC calculated using TRF time courses from 0 s to 0.4 s. Bars reflect the mean. Dots reflect data from individual participants. Right: Between-participant reliability for EEG reconstruction accuracy. Note that between-participant ICC is a measure across participants and thus does not have participant-unique data points. **B:** Generalizability across sessions (i.e., session 1 vs session 2 for DM stories): Same as in panel A.

Within-participant ICC for clear stories was 0.53 for younger and 0.54 for older adults (t_40_ = 0.087, p = 0.931, d = 0.027). Within-participant ICC for stories in babble was about 0.46 and 0.61 for younger and older adults, respectively (t_40_ = 1.839, p = 0.073, d = 0.568; Figure 8A, middle).

Between-participant ICC for the EEG reconstruction accuracy for younger adults was about 0.7 and 0.25 for clear stories and stories in noise, respectively. For older adults, between-participant ICC was about 0.55 and 0.75 for clear stories and stories in noise, respectively (Figure 8A, right).

### Across-story generalizability: Between-sessions focus

DM stories were used to investigate the generalizability of neural responses across stories and sessions. That is, the same reliability analyses (using ICC) as for SM stories were calculated for DM stories. Time courses of between-participant ICC for TRF amplitudes are shown in Figure 8B (left), showing peak ICC values of about 0.5-0.6 (moderate) at around 0.1 s, although ICC for clear stories for younger adults was lower (<0.5).

The within-participant ICC was 0.48 for the clear story and the story in noise for younger adults (i.e., low reliability; Koo and Li, 2016). For older adults, within-participant ICC was 0.53 and 0.6 for the clear story and the story in noise, respectively. There were no significant differences between speech-clarity conditions or age groups (ps > 0.15; Figure 8B, middle).

Between-participant ICC for the EEG reconstruction accuracy was about 0.6 for older adults under clear and noise conditions, whereas, for younger adults, it was about 0.15 and 0.6 for clear stories and stories in noise, respectively (Figure 8B, right).

### Across-story generalizability: Between-sessions, same speaker focus

DB stories were used to investigate generalizability across sessions for stories that mirror an audiobook and were spoken by the same speaker. Neural responses are shown in Figure 9. The rmANOVA for the 0.03-0.06 s time window revealed larger TRF amplitudes for older compared to younger adults (F_1,42_ = 38.877, p = 1.8 · 10^-7^, ω^2^ = 0.306; Figure 9B), whereas the effect of Session and the Session × Age Group interaction were not significant (ps > 0.5). The rmANOVA for the 0.09-0.13 s time window revealed no effects (ps > 0.25). The rmANOVA for reconstruction accuracy revealed larger correlations for older compared to younger adults (F_1,42_ = 15.868, p = 2.6 10^-4^, ω^2^ = 0.147; Figure 9C), whereas the effect of Session and the Session × Age Group interaction were not significant (ps > 0.05).

**Figure 9:**
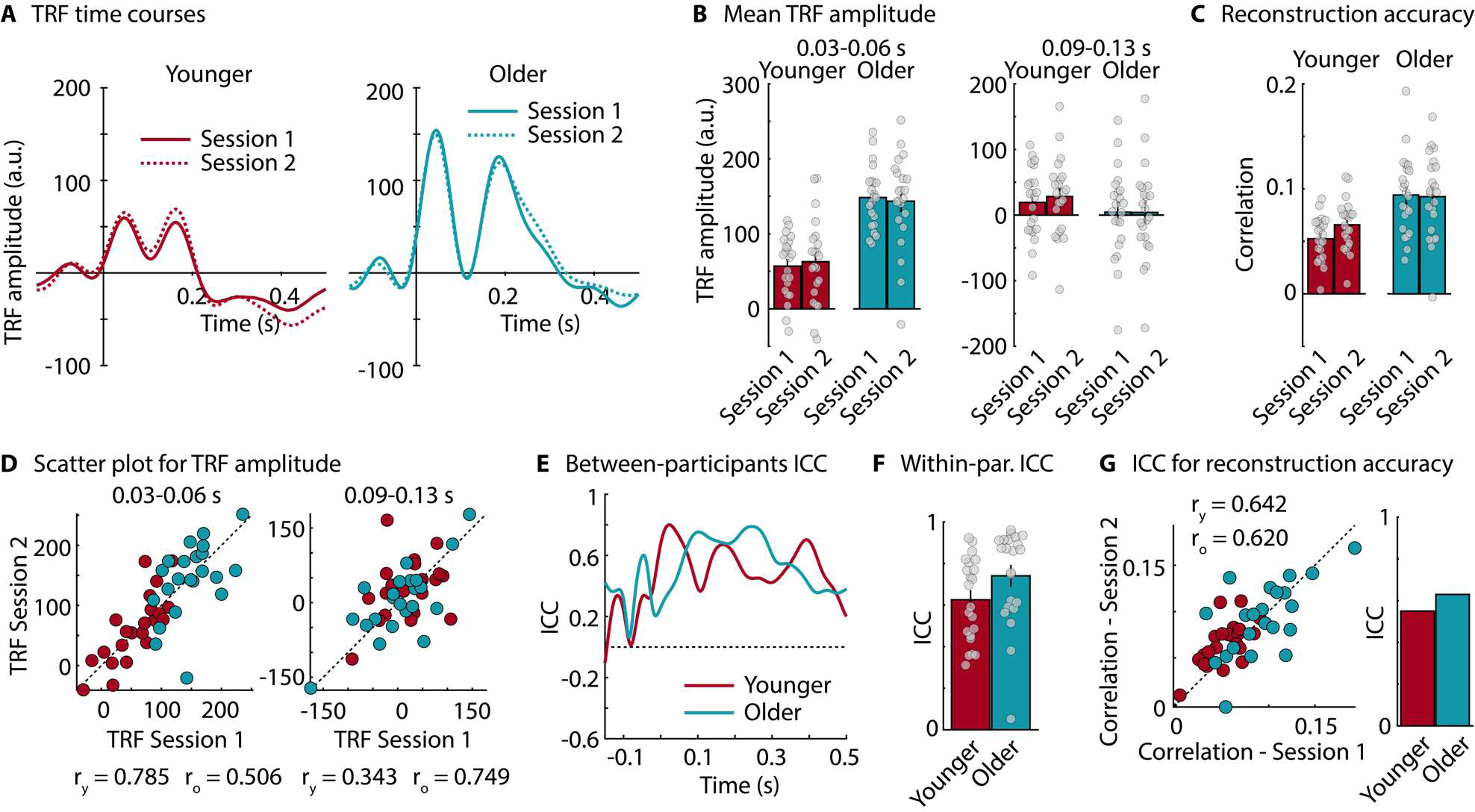
Neural responses and intra-class correlation for DM stories. **A:** Temporal response functions (TRFs). **B:** Mean TRF amplitude for the 0.03-0.06 s and the 0.09-0.13 s time windows. **C:** EEG reconstruction accuracy. **D:** 45-degree scatter plots for the TRF amplitude in the 0.03-0.06 s and the 0.09-0.13 s time windows. Pearson correlations are provided below each plot. The subscripts y and o indicate correlations for younger and older adults, respectively. **E:** Between-participant ICC calculated for 0.05 s sliding time windows centered on each time point. **F:** Within-participant ICC calculated using TRF time courses from 0 s to 0.4 s. **G:** 45-degree scatter plot and ICC for reconstruction accuracy. Pearson correlations are provided in the scatter plot. The subscripts y and o indicate correlations for younger and older adults, respectively. Dots in the different panels reflect data from individual participants.

Scatter plots of TRF amplitudes are shown in Figure 9D. The time courses of between-participant ICC for TRF amplitudes in Figure 9E show ICC values of about 0.5-0.7 (moderate) between 0 and 0.2 s, although ICC for younger adults was lower around 0.1 s (<0.5). Within-participant ICC was 0.62 and 0.74 for younger and older adults, respectively (i.e., moderate; Koo and Li, 2016; Figure 9F) and there was no difference between age groups (p > 0.05). Between-participant ICC for the EEG reconstruction accuracy was about 0.55 and 6.3 for younger and older adults, respectively (Figure 9G).

### Comparing reliability and generalizability

We also assessed whether within-participant ICC values differed between assessment types: ERP reliability (session 1 vs session 2), TRF reliability (SM stories, session 1 vs session 2), within-session TRF generalizability across story/speaker (SM vs DM stories, session 1), TRF generalizability across session/story/speaker (DM stories, session 1 vs session 2), TRF generalizability across session/story (DB stories, session 1 vs session 2).

The rmANOVA revealed a significant main effect of Assessment Type (F_4,160_ = 22.958, p = 5.2 · 10^-15^, ω^2^ = 0.219; Figure 10). Post hoc comparisons showed that ICC for ERP reliability was greater than ICC for TRF reliability (t_41_ = 6.012, p_Holm_ = 9.6 · 10^-8^, d = 1.062), within-session TRF generalizability across story/speaker (t_41_ = 7.760, p_Holm_ = 8.5 · 10^-12^, d = 1.371), TRF generalizability across session/story/speaker (t_41_ = 8.516, p_Holm_ = 1.1 · 10^-13^, d = 1.505), and TRF generalizability across session/story (t_41_ = 4.337, p_Holm_ = 1.8 · 10^-4^, d = 0.766). ICC for TRF generalizability across session/story (DB stories) was also greater than ICC for within-session TRF generalizability across story/speaker (t_41_ = 3.423, p_Holm_ = 0.004, d = 0.605) and TRF generalizability across session/story/speaker (t_41_ = 4.179, p_Holm_ = 2.8 · 10^-4^, d = 0.738). No other effects, including age effects, were significant (ps > 0.05).

**Figure 10:**
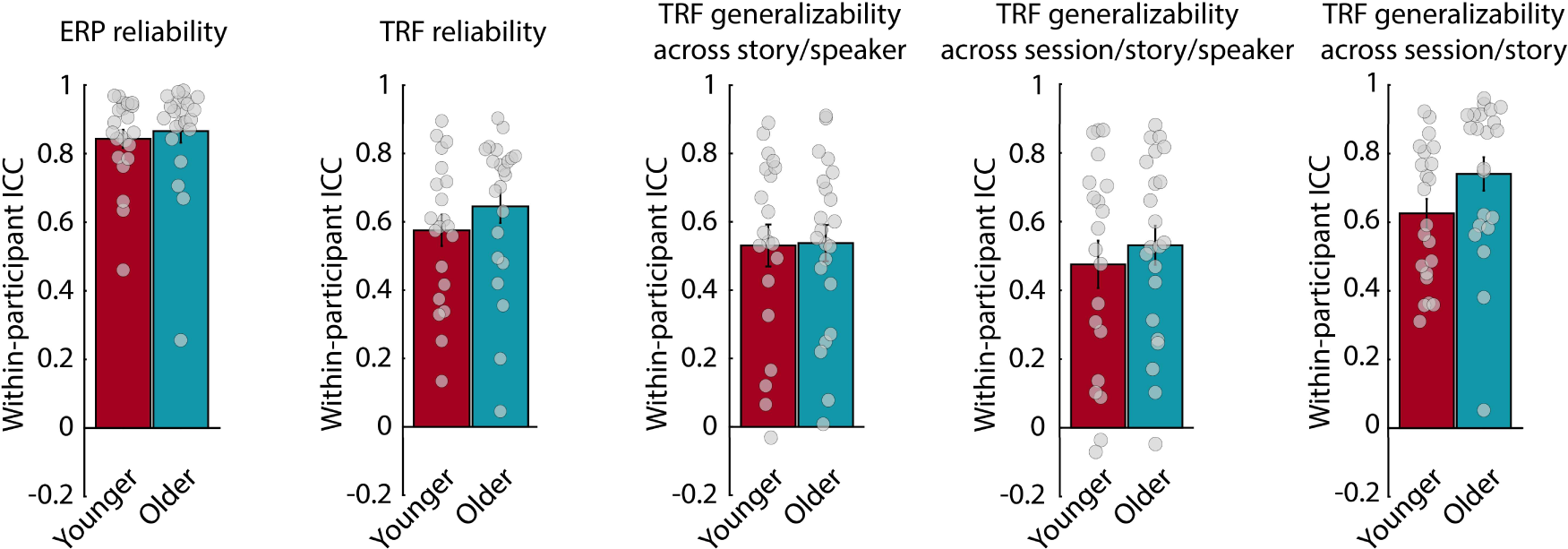
Comparison of reliability and generalizability. Within-participant ICC using TRF time courses from 0 s to 0.4 s. ICC is shown for ERP reliability (responses to noise bursts; session 1 vs session 2), TRF reliability (responses to the clear SM story; session 1 vs session 2), within-session TRF generalizability across story/speaker (responses to clear stories in session 1; SM vs DM), TRF generalizability across session/story/speaker (responses to clear DM stories; session 1 vs session 2), TRF generalizability across session/story (responses to DB stories; session 1 vs session 2). Bar graphs show the mean ICC. Error bars reflect the standard error of the mean. Dots reflect data from individual participants.

## Discussion

Assessment of the neural tracking of continuous, naturalistic speech is increasingly used to understand speech encoding and considered a potential clinical biomarker, for example, for age-related hearing loss. However, to use neural speech tracking as a biomarker requires knowledge about its reliability. The current study investigated the reliability and generalizability of neural speech tracking in younger and older adults while they listened to stories and EEG was recorded in two separate sessions. Neural responses to noise bursts were used as a benchmark for which we expected high reliability. Early responses to noise bursts (∼0.05 and ∼0.1 s), neural-speech tracking responses ∼0.05 s (P1), and EEG reconstruction accuracy were larger for older compared to younger adults. Critically, reliability of neural speech tracking was moderate (ICC ∼0.5-0.75) in younger and older adults, and there was a tendency for reliability to be larger in older adults for speech presented in moderate background babble. Reliability for responses to noise bursts was higher (ICC >0.8) than the speech-tracking reliability in both younger and older adults. Neural-speech tracking responses also moderately generalized across different stories (ICC ∼0.5-0.6). Overall, the current study provides an important step in the development of an objective marker of speech encoding that can be used in clinical contexts.

### Age-related enhancement of neural responses to white noise and stories

We observed larger responses for older compared to younger adults, for both noise bursts and spoken stories. This response enhancement has been observed frequently in older relative to younger adults for speech and non-speech sounds (Tremblay et al., 2003; Alain et al., 2012; Herrmann et al., 2013; Alain et al., 2014; Bidelman et al., 2014; Herrmann et al., 2016; Presacco et al., 2016b; Millman et al., 2017; Brodbeck et al., 2018; Decruy et al., 2019; Harris et al., 2022), and is sometimes more prominent for the earlier (∼0.05 s) than the later (∼0.1 s) cortical response (Alain et al., 2014; Harris et al., 2022; Figures 5 and 6).

A variety of possible mechanisms for the age-related response enhancement have been discussed, including a loss of cortical inhibition resulting from peripheral deafferentation (Auerbach et al., 2014; Chambers et al., 2016; Resnik and Polley, 2017; Salvi et al., 2017; Herrmann and Butler, 2021b), recruitment of additional cortical resources (Brodbeck et al., 2018; Gillis et al., 2022), and increased attention or effort (Vanthornhout et al., 2019; Decruy et al., 2020b; Gillis et al., 2022). However, cognitive factors are unlikely the sole contributors to the age-related response enhancement, given that it is present also under distracted listening conditions (Herrmann et al., 2016; Herrmann et al., 2018; Harris et al., 2022; Figure 3).

### Reliability of neural speech tracking

We investigated the reliability of neural speech tracking by recording EEG from individuals while they listened to the same stories twice in separate sessions. Younger and older adults experienced lower listening effort for the masked story in session 2 compared to session 1, indicating that prior knowledge about speech can reduce listening effort (Pichora-Fuller et al., 1995; Obleser and Kotz, 2010; Signoret et al., 2011; Holmes et al., 2018; Herrmann and Johnsrude, 2020). However, younger adults found listening to the same story a second time less enjoyable and absorbing than when they listened to it for the first time, whereas older adults found them similarly enjoyable and absorbing both times (Figure 4). Our observation that listening effort, enjoyment, and absorption are reduced when listening to the same story several times perhaps suggests that different stories should be used in clinical contexts when individuals are assessed several times.

There were no differences in TRF amplitude or EEG reconstruction accuracy between session 1 and 2, and peak reliability of neural-tracking responses (TRF, accuracy) was moderate for both age groups (ICC ∼0.5-0.75; Koo and Li, 2016; Figure 7), although it appeared that in several analyses reliability was lower somewhat lower in younger than older adults (e.g., for EEG reconstruction accuracy, Figure 7C). This may, in part, be due to larger responses for older compared to younger adults, but might also be related to the decrease in absorption and enjoyment from session 1 to session 2 for younger adults that was absent in older adults. The reliability for neural speech tracking was lower compared to the reliability for responses to noise bursts (showing good reliability; ICC >0.75; Koo and Li, 2016), which is consistent with the good reliability for simple sound stimuli observed previously (Tervaniemi et al., 1999; Legget et al., 2017; Bidelman et al., 2018).

Neural speech tracking is increasingly used to investigate clinical phenomena, such as hearing loss (Presacco et al., 2019; Decruy et al., 2020a; Schmitt et al., 2022), and researchers have suggested that neural speech tracking could be an important biomarker (Gillis et al., 2022; Palana et al., 2022; Schmitt et al., 2022). However, a reliability of 0.7 or higher has been recommended for measures used in clinical research (Nunnally and Bernstein, 1994; Frost et al., 2007; Mokkink et al., 2022) and it thus appears that the moderate (ICC ∼0.5-0.75) reliability for neural speech tracking in older adults may not be sufficiently high to meet this criterion, or only in specific time windows for older adults.

The current data further help quantify the upper bound of how well neural speech tracking can correlate with or predict a clinical condition. That is, the maximum correlation between two measures is equal to the square root of the product of their reliabilities (sqrt(reliability of Measure A × reliability of Measure B); Bedeian et al., 1997; Goodwin and Leech, 2006; Bedeian, 2014). Standard clinical measures tend to have good test-retest reliability (ICC ∼0.8; sometimes correlation instead of ICC is provided). For example, good-to-high reliability has been observed for audiometric assessments (McClannahan et al., 2021) and cognitive assessments (Montreal Cognitive Assessment; Nasreddine et al., 2005; Gupta et al., 2019; Lee et al., 2022). Reliability values in the current study were somewhat variable across neural-tracking measures, particularly, in younger adults and for reconstruction accuracy. Nevertheless, assuming a moderate ∼0.6 reliability of the neural-tracking response and a reliability of 0.8 for audiological assessments, we would expect a maximum correlation of 0.69 between the two measures.

Critically, the current reliability assessment has focused on the neural tracking of the speech envelope. The speech envelope has been used most (Lalor and Foxe, 2010; Ding and Simon, 2012; Hertrich et al., 2012; Ding et al., 2015; Fiedler et al., 2019; Fiedler et al., 2021), can be easily calculated, and may thus be particularly useful as a clinical biomarker compared to recent approaches to assess tracking of semantic features of speech that require more complex analyses (Broderick et al., 2018; Broderick et al., 2021; Marlies et al., 2021; Yasmin et al., 2023). Nevertheless, future work should further investigate the reliability of the neural tracking of linguistic features of speech.

### Generalizability of neural speech tracking across stories

Diagnosis or treatment of a clinical condition can involve repeated assessments of a person using the same measure or procedure, for example, when evaluating intervention progress for the treatment of hearing loss. A biomarker of speech processing should thus be independent of specific speech stimuli and instead generalize across stimuli to avoid prior knowledge affecting the measurement outcome. The current generalizability data show that across-story ICC did not differ from same-story ICC (strict reliability), although the former was numerically lower (Figure 10). While ICC values for reliability (same story) and generalizability (different stories) were only moderate, there seems to be little indication that using neural speech tracking as an assessment tool would suffer from using different stories in the case that two or more assessments are needed.

Critically, our results suggest that generalizability may be highest if audiobooks (here story-book stories) spoken by the same person are used as speech materials (Figure 9 and 10). The Moth stories are highly engaging, enjoyable to participants, and reflect real-life speech with pauses, disfluencies, and other idiosyncrasies. Some work suggests The Moth stories are more enjoyable and absorbing than story-book stories (Mathiesen et al., 2023), although this difference was not observed in the current study. We recommend that clinical research could perhaps rely on the more controlled audiobooks spoken by the same person if repeated assessments are needed, while enjoyment for such stories, relative to more real-life speech, is likely reduced only minimally.

## Conclusions

The current study investigated the reliability and generalizability of neural speech tracking in younger and older adults. Participants listened to stories that either repeated in two sessions (to test reliability) or differed across sessions (to test generalizability). We observed larger neural responses for older compared to younger adults during story listening, consistent with a loss of inhibition in the aged auditory system. Reliability for the neural-tracking response was moderate (ICC ∼0.5-0.75) and somewhat lower and more variable across story and speech-clarity conditions in younger compared to older adults. Generalizability across stories appeared greatest when audiobook-like stories spoken by the same person were used as speech materials. The current data provide results critical for the development of an objective biomarker of speech processing, but also suggest that further work is needed to increase the reliability of the neural-tracking response to meet clinical standards.

## Acknowledgements

We thank Christie Tsagopoulos and Tazeen Atif for their help with data collection. The research was supported by the Canada Research Chair program (CRC-2019-00156, 232733) and the Natural Sciences and Engineering Research Council of Canada (Discovery Grant: RGPIN-2021-02602).

## Authors contributions

**Ryan A. Panela:** Formal analysis, visualization, writing – original draft, writing - review and editing. **Francesca Copelli:** Methodology, formal analysis, data curation, writing - review and editing. **Björn Herrmann:** Conceptualization, methodology, formal analysis, investigation, data curation, writing - original draft, writing - review and editing, visualization, supervision, project administration, funding acquisition.

